# Parallel and scalable workflow for the analysis of Oxford Nanopore direct RNA sequencing datasets

**DOI:** 10.1101/818336

**Authors:** Luca Cozzuto, Huanle Liu, Leszek P. Pryszcz, Toni Hermoso Pulido, Julia Ponomarenko, Eva Maria Novoa

## Abstract

The direct RNA sequencing platform offered by Oxford Nanopore Technologies allows for direct measurement of RNA molecules without the need of conversion to complementary DNA, fragmentation or amplification. As such, it is virtually capable of detecting any given RNA modification present in the molecule that is being sequenced, as well as provide polyA tail length estimations at the level of individual RNA molecules. Although this technology has been publicly available since 2017, the complexity of the raw Nanopore data, together with the lack of systematic and reproducible pipelines, have greatly hindered the access of this technology to the general user. Here we address this problem by providing a fully benchmarked workflow for the analysis of direct RNA sequencing reads, termed *MasterOfPores*. The pipeline converts raw current intensities into multiple types of processed data, providing metrics of the quality of the run, quality-filtering, base-calling and mapping. The output of the pipeline can in turn be used to compute per-gene counts, RNA modifications, and prediction of polyA tail length and RNA isoforms. The software is written using the NextFlow framework for parallelization and portability, and relies on Linux containers such as Docker and Singularity for achieving better reproducibility. The *MasterOfPores* workflow can be executed on any Unix-compatible OS on a computer, cluster or cloud without the need of installing any additional software or dependencies, and is freely available in Github (https://github.com/biocorecrg/master_of_pores). This workflow will significantly simplify the analysis of nanopore direct RNA sequencing data by non-bioinformatics experts, thus boosting the understanding of the (epi)transcriptome with single molecule resolution.

## INTRODUCTION

Next generation sequencing (NGS) technologies have revolutionized our understanding of the cell and its biology. However, NGS technologies are heavily limited by their inability to sequence long reads, thus requiring complex bioinformatic algorithms to assemble back the DNA pieces into a full genome or transcriptome. Moreover, NGS technologies require a PCR amplification step, and as such, they are typically blind to DNA or RNA modifications (Novoa et al., 2017).

The field of epitranscriptomics, which studies the biological role of RNA modifications, has experienced an exponential growth in the last few years. Systematic efforts coupling antibody immunoprecipitation or chemical treatment with next-generation sequencing (NGS) have revealed that RNA modifications are much more widespread than originally thought, are reversible (Jia et al., 2011), and can play major regulatory roles in determining cellular fate (Batista et al., 2014), differentiation (Furlan et al., 2019; Lee et al., 2019; Lin et al., 2017) and sex determination (Haussmann et al., 2016; Kan et al., 2017; Lence et al., 2016), among others. However, the lack of selective antibodies and/or chemical treatments that are specific for a given modification have largely hindered our understanding of this pivotal regulatory layer, limiting our ability to produce genome-wide maps for 95% of the currently known RNA modifications (Jonkhout et al., 2017).

Third-generation sequencing (TGS) platforms, such as the one offered by Oxford Nanopore Technologies (ONT), allow for direct measurement of both DNA and RNA molecules without prior fragmentation or amplification (Brown and Clarke, 2016), thus putting no limit on the length of DNA or RNA molecule that can be sequenced. In the past few years, ONT technology has revolutionized the fields of genomics and (epi)transcriptomics, by showing its wide range of applications in genome assembly (Jain et al., 2018), study of structural variations within genomes (Cretu Stancu et al., 2017), 3’ poly(A) tail length estimation (Krause et al., 2019), accurate transcriptome profiling (Bolisetty et al., 2015), identification of novel isoforms (Byrne et al., 2017; Križanovic et al., 2018) and direct identification of DNA and RNA modifications (Carlsen et al., 2014; Garalde et al.; Liu et al., 2019; Simpson et al., 2017). Thus, not only this technology overcomes many of the limitations of short-read sequencing, but importantly, it also can directly measure RNA and DNA modifications in their native molecules. Although ONT can potentially address many problems that NGS technologies cannot, the lack of proper standardized pipelines for the analysis of ONT output is greatly limited its reach to the scientific community.

To overcome these limitations, workflow management systems together with Linux containers offer an efficient solution to analyze large-scale datasets in a highly reproducible, scalable and parallelizable manner. In the last year, several workflows to analyze nanopore data have become available; however, these are mainly limited to genome assembly (e.g. Katuali; https://github.com/nanoporetech/katuali) and genome annotation (e.g. Pinfish; https://github.com/nanoporetech/pipeline-pinfish-analysis). Here we provide a scalable and parallelizable workflow for the analysis of direct RNA (dRNA) sequencing datasets, termed *MasterOfPores* (https://biocorecrg.github.io/master_of_pores/), which uses as input raw direct RNA sequencing FAST5 reads, and is aimed to facilitate the analysis of (epi)transcriptomics sequencing data. Specifically, *MasterOrPores* takes as input the raw fast5 reads produced by the sequencer, which can be either in single FAST5 of multi-FAST5 format, and performs quality control, filtering, and base-calling. For each step, it extracts metrics which are compiled in a final HTML report that can be easily visualized and analyzed by non-expert bioinformaticians. A direct RNA sequencing run produced by MinION or GridION devices, which typically comprises about 1M reads, takes ∼2 hours to analyze on a cluster using 100 nodes, each one with 8 CPUs, and ∼1 hour or less on a single GPU (see **Table 1** for detailed metrics). Moreover, the pipeline can also be run on the cloud (see section “Running on AWS”).

**Table 1.**
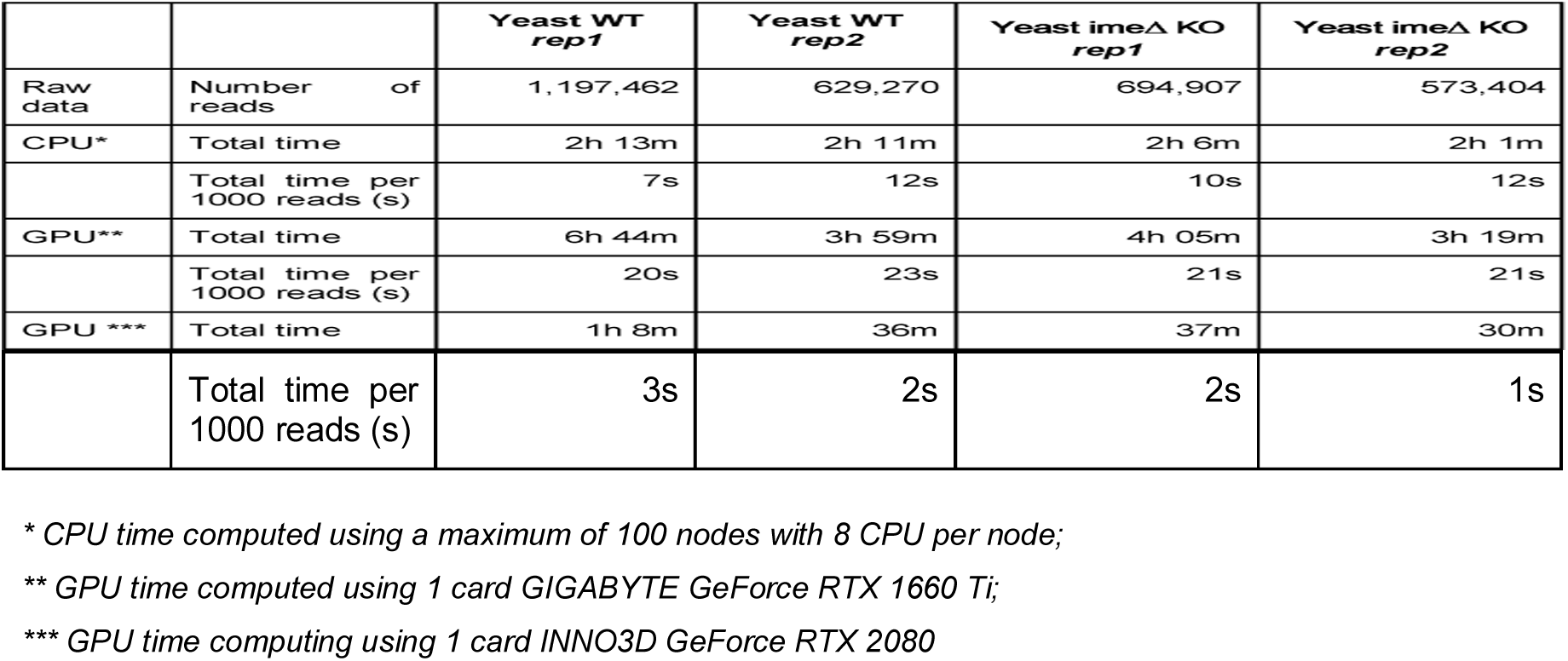
Comparison of computing time and RAM used to run the pipeline for the four *S. cerevisiae* polyA(+) direct RNA sequencing datasets used in this study, including subsets of one of the runs.

*MasterOfPores* simplifies the analysis of direct RNA sequencing data by providing a containerised pipeline implemented in the NextFlow framework. It is important to note that this approach avoids the heavy-lifting of installing dependencies by the user, and thus, is simple and accessible to any researcher without bioinformatics expertise. We expect that our workflow will greatly facilitate the access of Nanopore direct RNA sequencing to the community.

## RESULTS

### Overview of the *MasterOfPores* workflow

Workflow management systems together with Linux containers offer a solution to efficiently analyse large scale datasets in a highly reproducible, scalable and parallelizable manner. During the last years an increasing interest in the field has led to the development of different programs such as Snakemake (Köster and Rahmann, 2012), NextFlow (Di Tommaso et al., 2017), Galaxy (Afgan et al., 2018), SciPipe (Lampa et al., 2019) or GenPipes (Bourgey et al., 2019), among others. These tools enable the prototyping and deployment of pipelines by abstracting computational processes and representing pipelines as directed graphs, in which nodes represent tasks to be executed and edges represent either data flow or execution dependencies between different tasks.

Here we chose the workflow framework NextFlow (Di Tommaso et al., 2017) because of its native support of different batch schedulers (SGE, LSF, SLURM, PBS and HTCondor), cloud platforms (Kubernetes, Amazon AWS and Google Cloud) and GPU computing, which is crucial for processing huge volumes of data produced by nanopore sequencers. NextFlow has tight integration with lightweight Linux containers, such as Docker and Singularity. Automatic organization of intermediate results produced during the NextFlow pipeline execution allows reducing the complexity of intermediary file names and the possibility of name clashing. Continuous check-pointing with the possibility of resuming failed executions, interoperability and meticulous monitoring and reporting of resource usage are among other thought-after features of NextFlow. The executables of the presented pipeline have been bundled within Docker images accessible at DockerHub that can be converted on the fly into a Singularity image, thus allowing the HPC usage.

The *MasterOfPores* workflow includes all steps needed to process raw FAST5 files produced by Nanopore direct RNA sequencing and executes the following steps, allowing users a choice among different algorithms (**Figure 1**, see also **Figure S1**):

**Figure 1.**
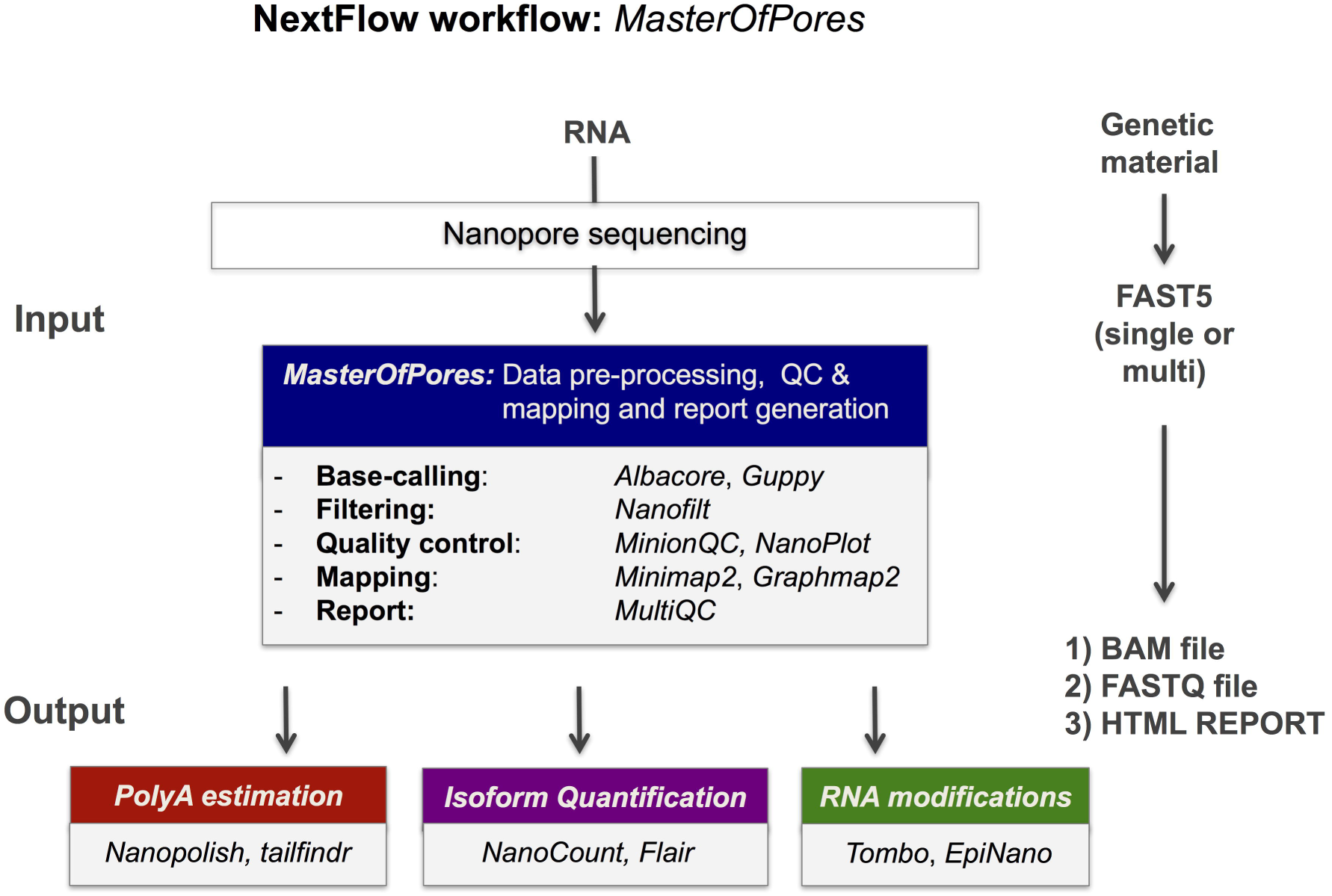
Overview of the *MasterOfPores* workflow for the processing of direct RNA nanopore sequencing datasets. The workflow accepts both single FAST5 and multi-FAST5 reads and includes 5 main steps: i) base-calling, i) filtering, iii) quality control, iv) mapping and v) final report building. The outputs generated by *MasterOfPores* (BAM file, fastq file, base-called fast5 files) can in turn be used as input to predict RNA modifications, polyA estimations, or per-transcript isoform quantifications. See also Figure S1.

i. Read base-calling with the algorithm of choice, using *Albacore* (https://nanoporetech.com) or *Guppy* (https://nanoporetech.com). This step can be run in parallel and the user can decide the number of files to be processed in a single job by using the command *--granularity*.
ii. Filtering of the resulting fastq files using Nanofilt (De Coster et al., 2018). This step is optional and can be run in parallel.
iii. Quality control of the base-called data using *MinIONQC* (Lanfear et al., 2019) and FastQC (http://www.bioinformatics.babraham.ac.uk/projects/fastqc).
iv. Read mapping to the reference genome or transcriptome using *minimap2* (https://github.com/lh3/minimap2) or *graphmap2* (https://github.com/lbcb-sci/graphmap2).
v. Quality control on the alignment using *NanoPlot* (https://github.com/wdecoster/NanoPlot) and *bam2stats* (https://github.com/lpryszcz/bin).
vi. Final report of the data processing using *multiQC* (https://github.com/ewels/MultiQC) that combines the single quality controls done previously, as well as global run statistics.

### Running *MasterOfPores:* installation, input, parameters and output

To run *MasterOfPores*, the following steps are required:

i. Install NextFlow (version 19.10.0)
  $ *curl -s https://get.nextflow.io | bash*
ii. Clone the MasterOfPores repository:
  $ *git clone --depth 1 biocorecrg/master_of_pores master_of_pores*
iii. Install Docker and/or Singularity (for Singularity, version 2.6.1 and Docker 19.03 or later are required): Docker: https://docs.docker.com/install/ Singularity: https://sylabs.io/guides/2.6/user-guide/quick_start.html#quick-installation-steps
iv. Download Nanopore base-calling algorithms: *guppy* with or without GPU support and or the albacore Wheel file (a standard built-package format used for Python distributions) and install them inside the *bin* folder inside the MasterOfPores directory. The users can place their preferred version of guppy and/or albacore in the *bin* folder (in the example below, albacore version 2.1.7 and guppy 3.1.5).
  $ *cd master_of_pores/bin*
  $ *tar -zvxf ont-guppy_3.1.5_linux64.tar.gz*
  $ *ln -s ont-guppy_3.1.5_linux64/ont-guppy/bin/guppy_*\*.
  $ *pip3 install --target=./albacore ont_albacore-2.1.7-cp36-cp36m-manylinux1_x86_64.whl*
  $ *ln -s albacore/bin/multi_to_single_fast5*
  $ *ln -s albacore/bin/read_fast5_basecaller.py*
v. Optional step: install CUDA drivers (only needed for GPU support): https://docs.nvidia.com/cuda/cuda-installation-guide-linux/index.html
vi. Run the pipeline (using singularity *or* docker):
  $ *nextflow run preprocessing.nf -with-singularity*
  $ *nextflow run preprocessing.nf -with-docker*

*MasterOfPores* can handle both single- and multi-FAST5 reads as input. To execute the workflow, several parameters can be defined by the user, including the choice of the basecaller software (albacore or guppy), choice of the mapper (minimap2 or graphmap2), as well as their command line options. If these are not specified by the user, the workflow will be run with default parameter settings (**Table 2**). The final report includes 4 different types of metrics: (i) *General statistics* of the input, including the total number of reads, GC content and number of identical base-called sequences; (ii) *Per-read statistics* of the input data, including scatterplots of the average read length versus sequence identity, the histogram of read lengths, and the correlation between read quality and identity; (iii) *Alignment statistics*, including the total number of mapped reads, the total number of mapped bases, the average length of mapped reads, and the mean sequence identity; (iv) *Quality filtering statistics*, including the number of filtered reads, median Q-score and read length, compared to those observed in all sequenced reads; and (v) *Per-read analysis of biases*, including information on duplicated reads, over-represented reads and possible adapter sequences (**Figure 2**).

**Table 2.**
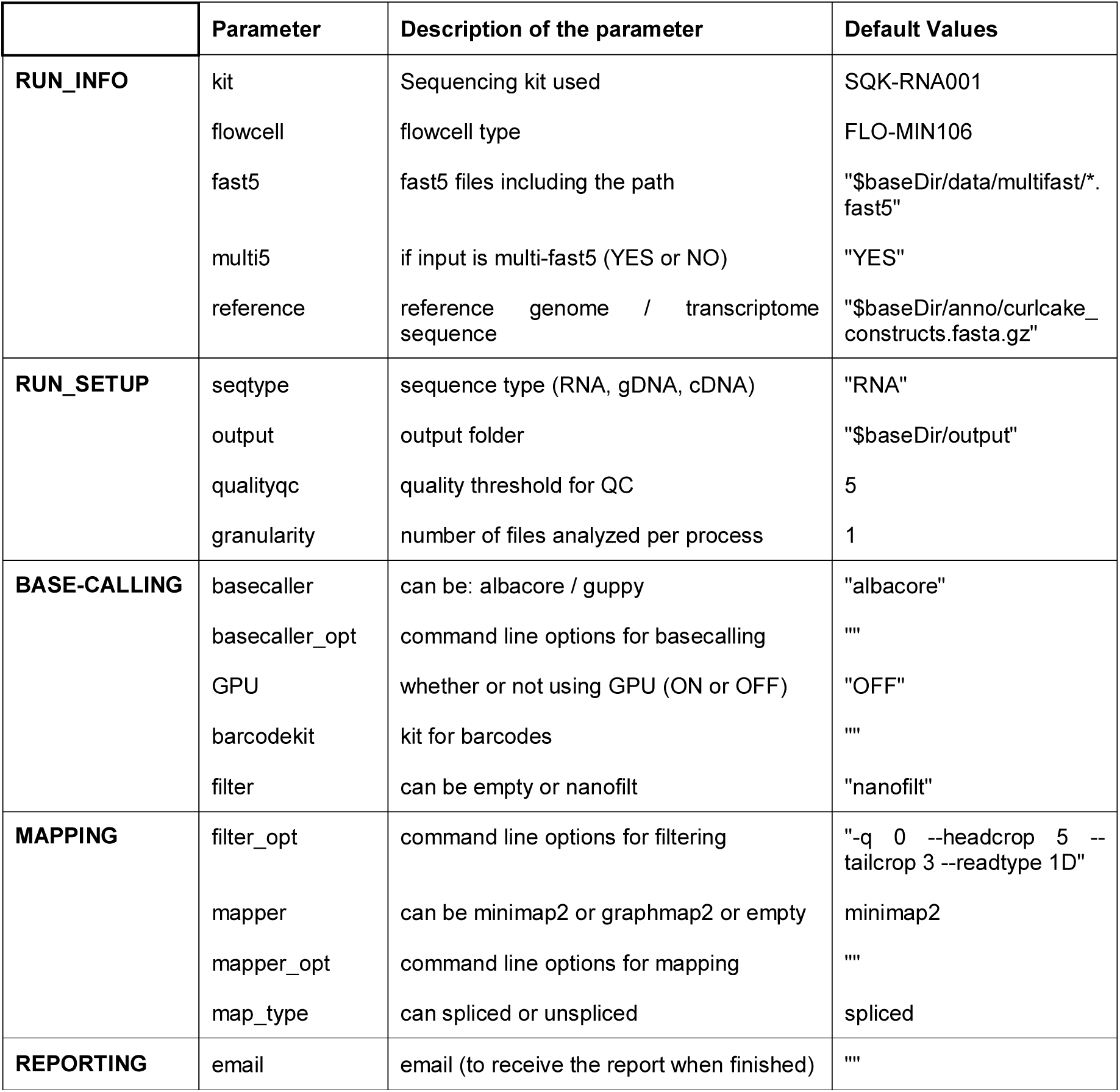
Settings and parameters that can be customized to run the MasterOfPores workflow.

**Figure 2.**
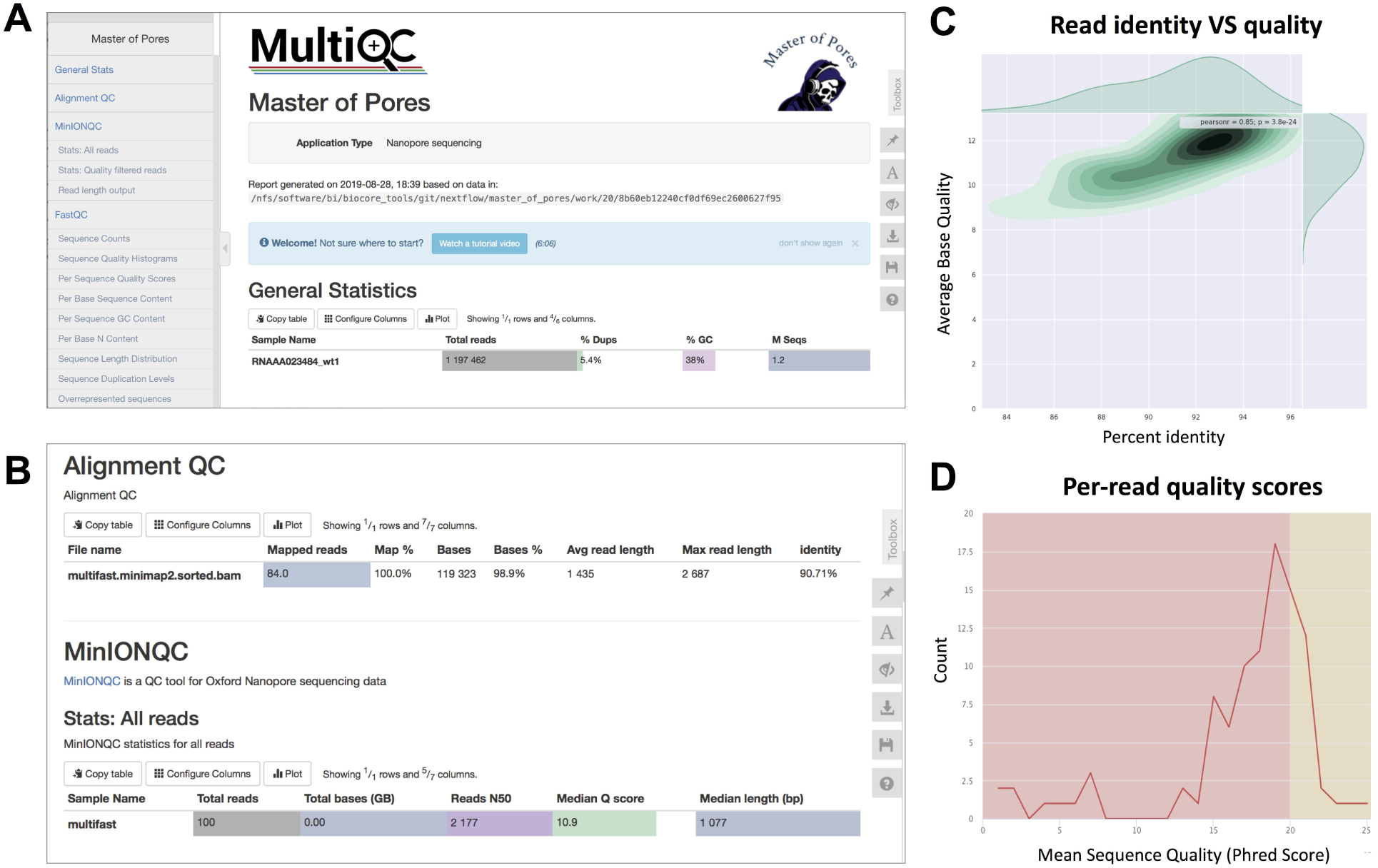
Snapshots of the final report generated by *MasterOfPores*. **(A)** Main menu and overview of the final report generated by *MasterOfPores*. **(B)** The report includes detailed metrics on the input reads (“MinIONQC”), as well as on the mapped reads (“AlignmentQC”). **(C**,**D)** Example of plots that are included as part of the *MasterOfPores* final report, some of which are generated by integrating Nanoplot (C) and FastQC (D) software.

The final outputs of the pipeline include:

- Basecalled fast5 files within the “fast5_files” folder.
- Filtered fastq files within “fastq_files” folder.
- QC reports within “QC” folder.
- Final report within “report” folder.
- Aligned reads in BAM files within the “aln” folder.

### Running *MasterOfPores* on the cloud (AWS Batch and AWS EC2)

Nanopore sequencing allows for real-time sequencing of samples. While GridION devices come with built-in GPUs that allows live base-calling, smaller MinION devices do not have built-in CPU or GPU. Thus, the user has to connect the MinION to a computer with sufficient CPU/GPU capabilities, or run base-calling after the sequencing. In all these contexts context, the possibility of running the *MasterOfPores* pipeline on the cloud presents a useful alternative.

The Amazon Web Services (AWS) Batch is a computing service that enables users to submit jobs to a cloud-based user-defined infrastructure, which can be easily set up via either code-based definitions or a web-based interface. Computation nodes can be allocated in advance or according to resource availability. Cloud infrastructure can be also deployed or dismantled on demand using automation tools, such as CloudFormation or Terraform.

Here we show that the *MasterOfPores* pipeline can be successfully implemented on the cloud, and provide the Terraform script for running *MasterOfPores* on the AWS Batch CPU environments, available in the GitHub repository (https://biocorecrg.github.io/master_of_pores/). To run the pipeline using the AWS Batch, the users only need to change a few parameters related to their accounts in a configuration file. The pipeline can be run from either a local workstation or an Amazon EC2 entrypoint instance initiated for this purpose (we recommend the latter). Data to be analysed can be uploaded to an Amazon S3 storage bucket.

Similarly, we also tested whether our pipeline could be run in Amazon Web Services (AWS) Elastic Compute Cloud (EC2), which is one of the most popular cloud services (**Table S1**). Compared to AWS Batch, to run any workflow in AWS EC2, the user must first create an Amazon Machine Image (AMI). The AMI can be created using the same instructions as provided in **File S1**, starting from the official Ubuntu 18.04 LTS AMI, and including both Docker and Singularity software with NVIDIA libraries support. Here we show that the resulting image can be used to run the *MasterOfPores* workflow with NVIDIA Tesla V100 GPU cards. Automation scripts to run *MasterOfPores* in AWS EC2 can be found in the GitHub repository (https://biocorecrg.github.io/master_of_pores/).

### Test case: Analysis of *Saccharomyces cerevisiae* SK1 polyA(+) RNA

#### Running the MasterOfPores pipeline on S. cerevisiae polyA(+) RNA

To benchmark the performance of the *MasterOfPores* workflow, we employed two publicly available direct RNA sequencing runs of polyA(+)-selected *S. cerevisiae* WT and ime4Δ strains, which had been sequenced using MinION and GridION devices, producing a total of ∼3 million reads (**Table 1**). We used up to 100 nodes with 8 CPUs for testing the base-calling in CPU mode and 1 node with 1 GPU card for testing the base-calling in GPU mode (**Table 1**).

The *MasterOfPores* pipeline was ran using guppy version 3.1.5 as the base-caller and minimap2 version 2.17 as the mapping algorithm. Reads were filtered by running nanofilt with the options “-q 0 -- headcrop 5 --tailcrop 3 --readtype 1D”. Filtered reads were mapped to the yeast SK1 fasta genome. Specifically, the command that was executed to run the pipeline with these settings was:

*$ nextflow run main.nf --basecaller guppy --multi5 YES --seqtype RNA \*

*--fast5 “FOLDERNAME/*\*.*fast5” --reference genome.fa.gz --mapper minimap2\*

*--filter nanofilt --filter_opt “-q 0 --headcrop 5 --tailcrop 3 --readtype 1D*”.

#### Benchmarking the time used for the analysis of S.cerevisiae polyA(+) RNA

Here we have tested the pipeline using both CPU and GPU computing. Specifically, we ran the pipeline on the following configuration: (i) a single CPU node (e.g., emulating the computing time on a single laptop); (ii) a CPU cluster with 100 nodes; (iii) a single mid-range GPU card (RTX2080); and (iv) a single high-end GPU card (GTX1080 Ti).

We found that the computing time required to run the pipeline on a single GPU card was significantly lower than the running time in parallel on a high-performance CPU cluster with 100 nodes, 8 cores per node (**Table 1**, see also **Table S1**). Moreover, we found that the computing time can be significantly reduced depending on the GPU card (base-calling step was ∼2X faster for GTX1080 Ti than for RTX2080).

#### Reporting resources used for the analysis of S. cerevisiae polyA(+) RNA

Taking advantage of the NextFlow reporting functions, the pipeline can produce detailed reports on the time and resources consumed by each process (**Figure 3**), in addition to the output files (bam, fastq) and final report (html), if the workflow is executed with parameters *-with-report* (formatted report) or*-with-trace* (plain text report). Running the base-calling on each multi-fast5 file in parallel on our dataset showed that the most memory intensive tasks (about 5 Gbytes) were the mapping step (using minimap2) and the quality control step (using Nanoplot) (**Table 3**), while the most CPU-intensive and time-consuming step (∼80min) was the base-calling (using Guppy) (**Table 4**).

**Table 3.**
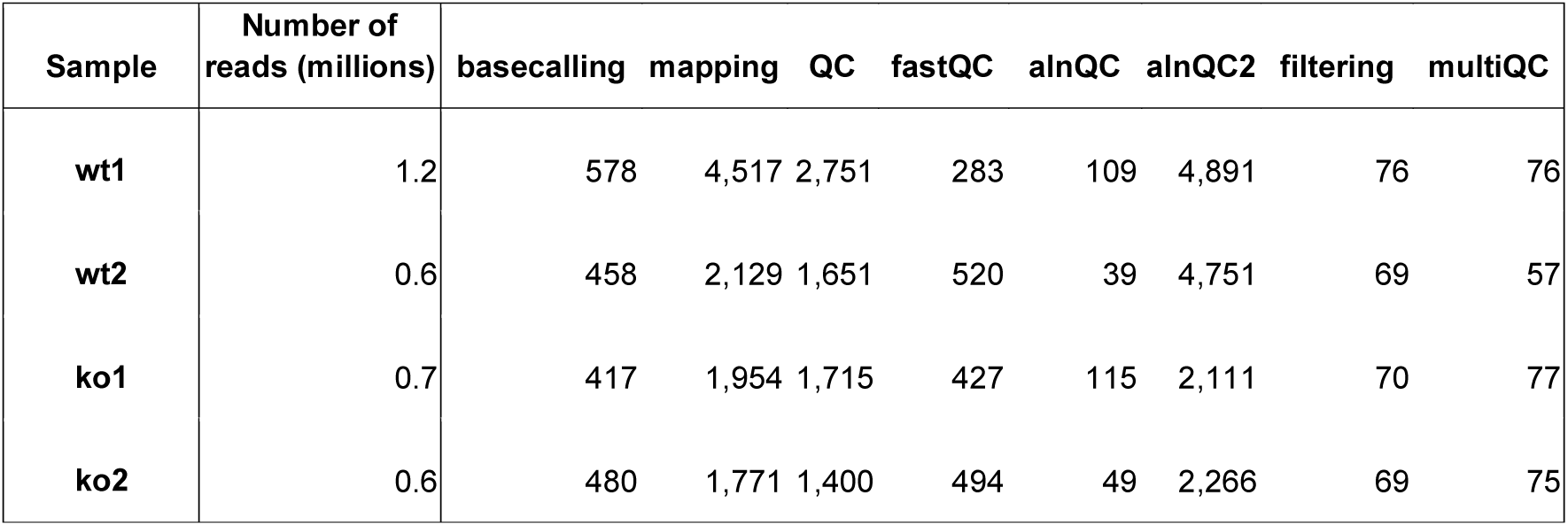
RAM peak (Mbytes) used by each of the pipeline modules.

**Table 4.**
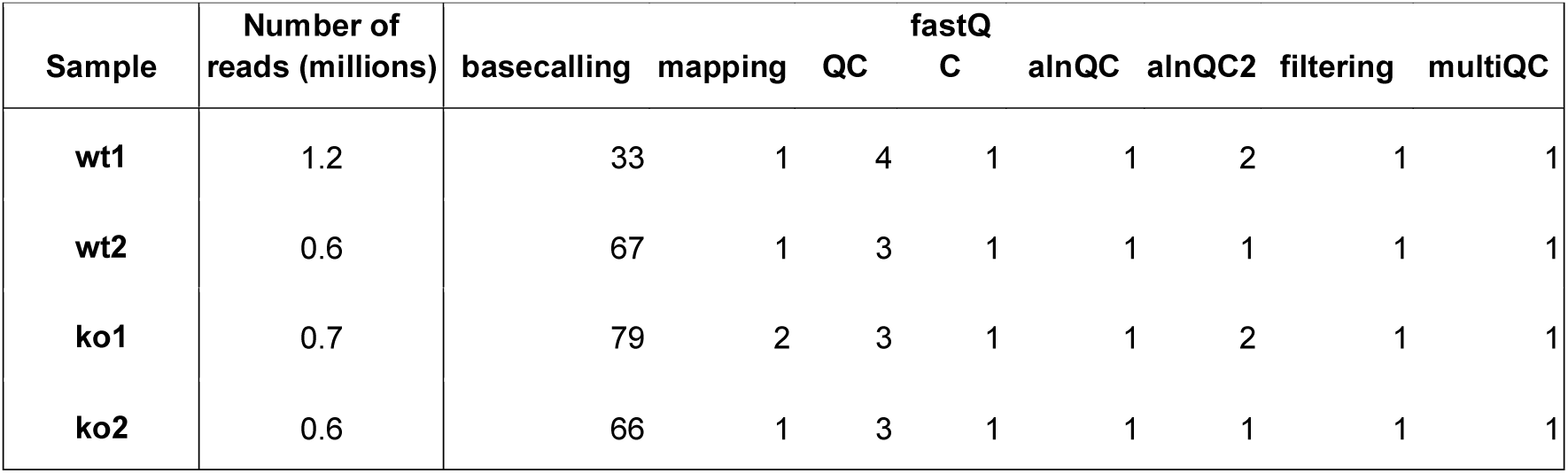
CPU time peak (min) used by each of the pipeline modules.

**Figure 3.**
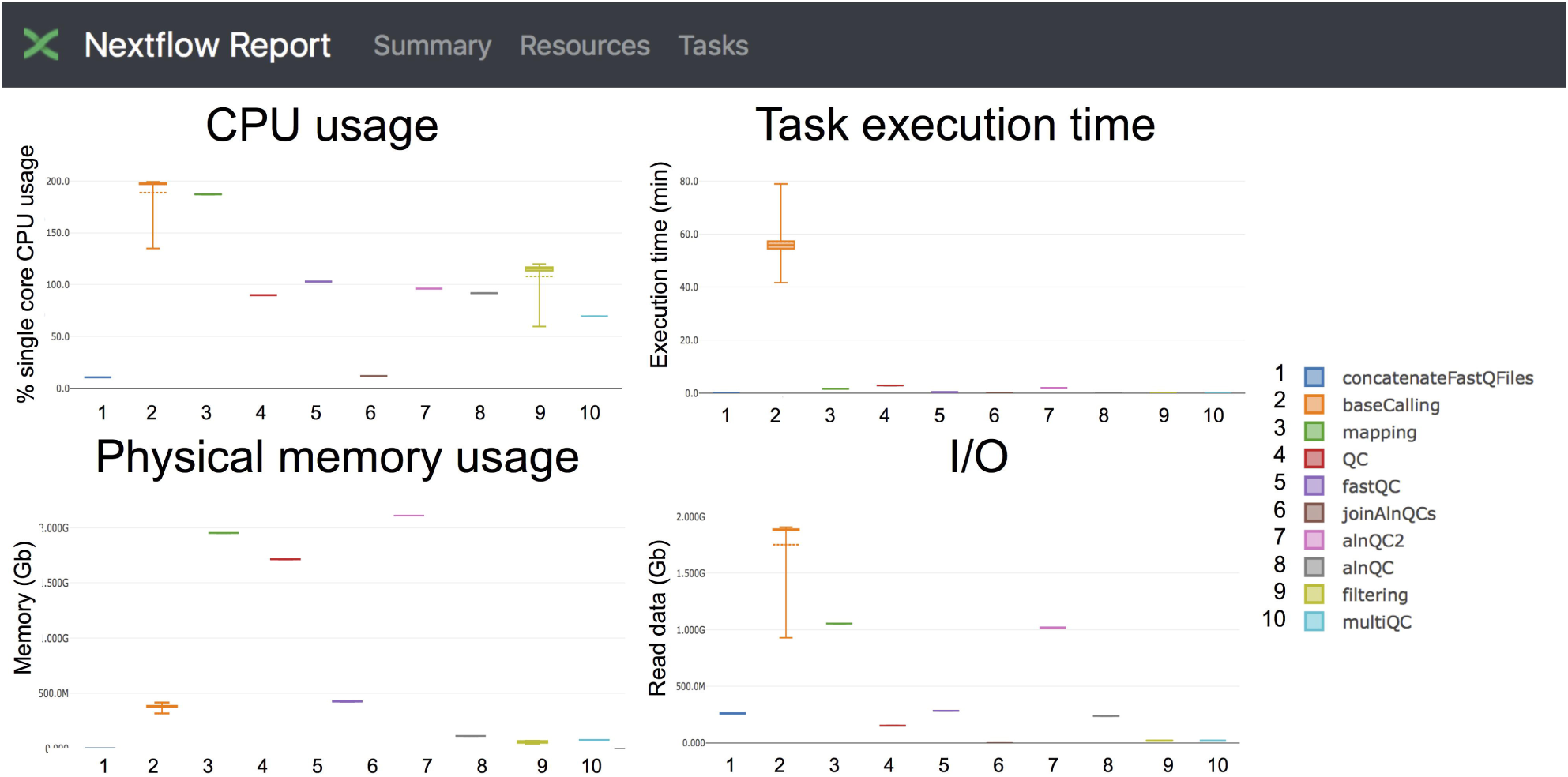
Snapshot of the NextFlow resources report. The report includes detailed information of the computing resources and time needed to execute each of the modules of the pipeline. Base-calling and mapping are the most CPU demanding tasks. The base-calling step is the longest to run, whereas mapping and generation of alignment QC metrics are the most memory-demanding tasks.

Finally, we should note that the latest (19.10.0) version of NextFlow allows the user to control the execution of a pipeline remotely. To enable this feature, the user needs to login to the https://tower.nf/ website developed by the NextFlow authors and retrieve a token for communicating with the pipeline. For doing that, the user should set this token as an environmental variable and run the pipeline as follows:

***$*** *export TOWER_ACCESS_TOKEN=YOUR_TOKEN*

*$ nextflow run main.nf -with-docker --granularity 300 -with-report -with-trace -bg -with-tower*

## DISCUSSION

The direct RNA sequencing technology offered by Oxford Nanopore technologies (ONT) offers the possibility of sequencing native RNA molecules, allowing to investigate the (epi)transcriptome at an unprecedented resolution, in full-length RNA molecules and in its native context. Although the direct RNA sequencing library preparation kit was made available in April 2017, only a modest number of researchers have started to adopt this new technology, partly due to the complexity of analyzing the resulting raw FAST5 data. Moreover, even in those cases when specific software and tools have been made available, the users typically experience many difficulties in installing dependencies and running the software. To overcome these issues and facilitate the data analysis of direct RNA sequencing to the general user, we propose the use of NextFlow workflows.

Specifically, we propose the use of *MasterOfPores* workflow for the analysis of direct RNA sequencing datasets, which is a containerised pipeline implemented in the NextFlow framework. *MasterOfPores* can handle both single- and multi-FAST5 reads as input, is highly customizable by the user (**Table 2**) and produces informative detailed reports on both the FAST5 data processing and analysis (MultiQC report, **Figure 2**) as well as on the computing resources used to perform each step (NextFlow report, see **Figure 3**). Thus, the current outputs of the *MasterOfPores* workflow include: (i) base-called FAST5 files, (ii) base-called fastq file, (iii) mapping BAM file, (iv) MultiQC report, and (v) NextFlow report. In the future we plan to integrate within the *MasterOfPores* workflow the software for the downstream analyses of direct RNA sequencing datasets including the PolyA tail length estimation, using Nanopolish (Workman et al., 2018) and tailfindr (Krause et al., 2019)) per-transcript isoform quantification and differential expression analysis, using Flair (Tang et al., 2018) and the analysis of RNA modifications, using Tombo (Stoiber et al., 2017) and EpiNano (Liu et al., 2019)).

The process of Nanopore read base-calling, that is, converting ion current changes into the sequence of RNA/DNA bases, has significantly improved during the last few years, mainly due to the adoption of deep learning approaches, such as the use of convolutional neural networks (CNNs) and recurrent neural networks (RNNs), which are currently the most commonly used strategies for base-calling. The adoption of RNN and CNN-based base-calling algorithms led to a dramatic improvement in base-calling accuracy. However, this came at the expense of a higher computational cost: only 5-10 reads can be base-called on 1 CPU core per second using the latest versions of the base-calling algorithms. The use of graphic processing units (GPUs) can greatly accelerate certain CPU-intensive computational tasks, thus allowing to process 50-500 reads per second (**Table S1**). We therefore developed our pipeline for both for CPU and GPU computing. Moreover, we provide the GPU-enabled docker image and detailed information on how to setup the GPU computing (see section: “Running MasterOfPores”). We encourage users to adopt the GPU computing for the analysis of Nanopore sequencing data whenever possible, as this option is both more time and cost-efficient.

## MATERIAL AND METHODS

### Code availability

The pipeline is publicly available at https://github.com/biocorecrg/master_of_pores under an MIT license. The example input data as well as expected outputs are included in the GitHub repository. Detailed information on program versions used can be found in the GitHub repository.

### Availability of Dockerfiles and Docker images

The pipeline uses software that is embedded within Docker containers. Dockerfiles are available in the GitHub repository (https://github.com/biocorecrg/master_of_pores/tree/master/docker/). The pipeline retrieves a specific Docker image from DockerHub, depending on whether the user requests to run the base-calling in GPU (https://cloud.docker.com/u/biocorecrg/repository/docker/biocorecrg/npbasecallgpu) or in CPU mode https://cloud.docker.com/u/biocorecrg/repository/docker/biocorecrg/npbasecallcpu) and it uses another one for the other tasks of the workflow (https://cloud.docker.com/u/biocorecrg/repository/docker/biocorecrg/nanopore).

### Integration of base-calling algorithms in the Docker images

Due to the terms and conditions that users agree to when purchasing Nanopore products, we are not allowed to distribute Nanopore software (binaries or in packaged form like docker images). While the original version of the *MasterOfPores* pipeline includes both guppy and albacore, we are not legally allowed to distribute it with the binaries. Therefore, here we only make available a version where the binaries must be downloaded and placed into a specific folder by the user. We expect future versions of *MasterOfPores* will include these softwares within the docker image once this issue is solved.

### CPU and GPU computing time and resources

The *MasterOfPores* workflow was tested both locally (using either CPU or GPU), as well as in the cloud (AWS). Computing times for each mode are shown in **Table 2**. CPU time was determined using a maximum of 100 nodes simultaneously with maximum 8 cores CPU per node (2.8-3.5 GHz, 80-130 Watt). GPU time was computed using either GIGABYTE GeForce RTX 1660 Ti (1536 CUDA cores @ 1770 MHz with 6GB of GDDR6 vRAM memory, 120 Watt) or INNO3D GeForce RTX 2080 (2944 CUDA cores @ 1710 MHz with 8 GB of GDDR6 vRAM memory, 225 Watt) or NVIDIA Tesla V100 (5120 CUDA cores + 640 Tensor cores @ 1462 MHz with 16 GB of HBM2 memory). For GPU computing, both system memory (RAM) and GPU memory (vRAM) are used. Base-calling with guppy typically uses 1 Gb or 4.2 Gb of vRAM in fast and high accuracy mode, respectively. As a result, only one base-calling process can be performed on above mentioned cards in high accuracy mode at given time. The execution time in the AWS EC2 p3.2xlarge instance involves reading files already placed in a previously set-up S3 storage bucket but not writing back output results into it.

### Data availability

Direct RNA sequencing datasets for *Saccharomyces cerevisiae SK1* PolyA(+) RNA were taken from publicly available GEO datasets (GSE126213)

## AUTHOR CONTRIBUTIONS

LC wrote the pipeline. HL optimized code and tested the pipeline. LP tested the pipeline and implemented GPU computing for containers. THP implemented and tested the workflow for AWS cloud computing. JP and EMN supervised the work. LC and EMN made figures and tables. LC, JP and EMN wrote the manuscript, with contributions from all authors.

## ACKNOWLEDGEMENTS

We thank all the members of the Novoa lab for their valuable insights and discussion. HL was supported by funds from the Australian Research Council (DP180103571 to EMN). LPP is supported by funding from the European Union’s H2020 research and innovation programme under Marie Sklodowska-Curie grant agreement No. 754422. This work was partly supported by the Spanish Ministry of Economy, Industry and Competitiveness (MEIC) (PGC2018-098152-A-100 to EMN). We acknowledge the support of the MEIC to the EMBL partnership, Centro de Excelencia Severo Ochoa and CERCA Programme / Generalitat de Catalunya. We would like to thank Benjamin Lang for letting us benchmark our pipeline on his hardware (GTX1080 Ti).

