## Supplementary information for "Parallel and scalable workflow for the analysis of Oxford Nanopore direct RNA sequencing datasets"

### SUPPLEMENTARY FIGURES

**Figure S1. Scheme of the individual steps, inputs and outputs of the *MasterOfPores* workflow.**

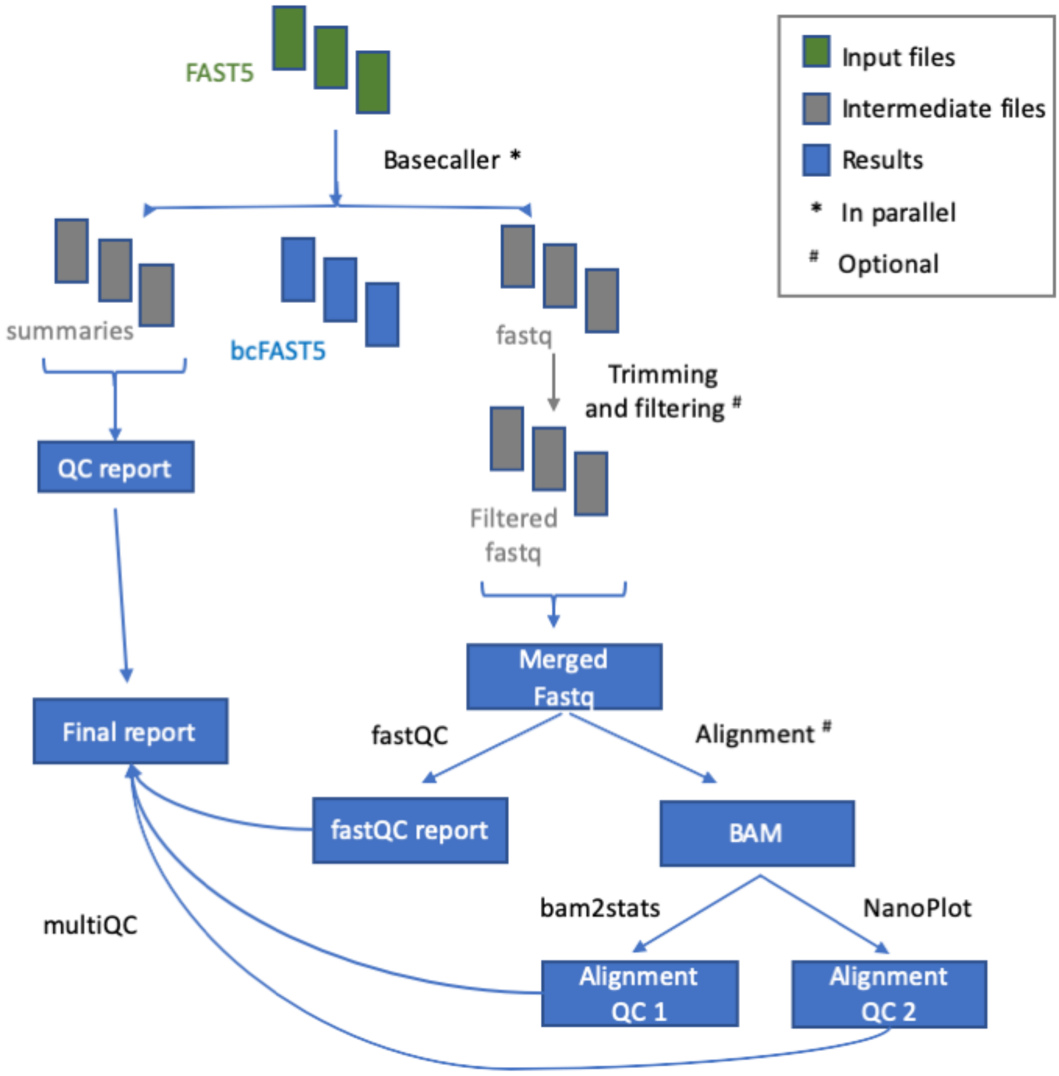

### SUPPLEMENTARY TABLES

**Table S1.** Scalability and resources (CPU or GPU computing time) used to run the workflow on increasing number of reads, benchmarked with the *S. cerevisiae* polyA(+) direct RNA sequencing WT dataset (replicate 1) used in this study.

| | | **Subset 1** | **Subset 2** | **Subset 3** | **Full dataset** |
|---|---|---|---|---|---|
| Raw data | Number of reads | 4,000 | 40,000 | 400,000 | 1,197,462 |
| CPU [a] | Total time | 16m | 2h 42m | 23h 34m | Not computed |
| | Total time per 1000 reads | 240s | 243s | 212s | Not computed |
| CPU [b] | Total time | 15m | 17m | 29m | 2h 13m |
| | Total time per 1000 reads | 225s | 22s | 4s | 6s |
| GPU [c] | Total time | 2m | 14m | 2h 15m | 6h 44 |
| | Total time per 1000 reads | 30s | 21s | 20s | 20s |
| GPU [d] | Total time | 45s | 3m | 21m | 1h 8m |
| | Total time per 1000 reads | 11s | 4s | 3s | 3s |
| GPU[e] | Total time | 2m | 4m | 34m | 1h44m |
| | Total time per 1000 reads | 30s | 6s | 5s | 5s |

[a] *CPU time computed using a maximum of 1 node with 8 CPU per node*

[b] *CPU time computed using a maximum of 100 nodes with 8 CPU per node;*

[c] *GPU time computed using 1 card GIGABYTE GeForce RTX 1660 Ti;*

[d] *GPU time computing using 1 card INNO3D GeForce RTX 2080*

[e] *GPU time computing using AWS p3.2xlarge instance (NVIDIA Tesla V100)*

### SUPPLEMENTARY FILES

#### File S1: Installation instructions for NVIDIA drivers, CUDA, Docker and Singularity

Instructions below have been tested with Ubuntu 18.04.

1. Install CUDA with the latest NVIDIA drivers

```
# install cuda with nvidia-drivers
sudo apt update && apt install -y software-properties-common headers-$(uname -r)
wget https://developer.download.nvidia.com/compute/cuda/repos/ubuntu1804/x86_64/cuda-ubuntu1804.pin
sudo mv cuda-ubuntu1804.pin /etc/apt/preferences.d/cuda-repository-pin-600
sudo apt-key adv --fetch-keys
https://developer.download.nvidia.com/compute/cuda/repos/ubuntu1804/x86_64/7fa2af80.pub
sudo add-apt-repository "deb
http://developer.download.nvidia.com/compute/cuda/repos/ubuntu1804/x86_64/ /"
sudo apt-get update && apt-get -y install cuda
```

2. Install Singularity v2.6.1

```
# install Singularity v2.6.1
sudo apt install libarchive-dev
export VERSION=2.6.1 && # adjust this as necessary \
wget https://github.com/sylabs/singularity/releases/download/${VERSION}/singularity-
${VERSION}.tar.gz
tar -xzf singularity-${VERSION}.tar.gz && \
cd singularity-${VERSION}
./configure -- prefix=/usr/local && make && sudo make install
```

3. Test Singularity installation and GPU-support

```
# Singularity running docker container without CUDA-toolkit
singularity exec --nv docker://ubuntu nvidia-smi
# Singularity running docker container with CUDA-toolkit installed
singularity exec --nv docker://nvidia/cuda nvidia-smi
```

4. Install the latest Docker-CE

```
# install docker-ce
sudo apt-get update && sudo apt-get install \
   apt-transport-https ca-certificates \
   curl gnupg-agent software-properties-common
curl -fsSL https://download.docker.com/linux/ubuntu/gpg | sudo apt-key add -
sudo apt-key fingerprint 0EBFCD88
sudo add-apt-repository \
  "deb [arch=amd64] https://download.docker.com/linux/ubuntu \
  $(lsb_release -cs) \
  stable"
sudo apt-get update && sudo apt-get install docker-ce docker-ce-cli containerd.io
# check if docker works
sudo docker run hello-world
# add your user to docker group - to run docker without sudo
sudo usermod -aG docker $USER
# and logout and login for the changes to start working
```

5. Install nvidia-docker for GPU-support in Docker

```
# install nvidia docker
distribution=$(. /etc/os-release;echo $ID$VERSION_ID)
curl -s -L https://nvidia.github.io/nvidia-docker/gpgkey | sudo apt-key add -
curl -s -L https://nvidia.github.io/nvidia-docker/$distribution/nvidia-docker.list | sudo tee
/etc/apt/sources.list.d/nvidia-docker.list
sudo apt-get update && sudo apt-get install -y nvidia-container-toolkit
sudo service docker restart
```

6. Test Docker installation

```
# run docker container with CUDA-toolkit installed
docker run -i --gpus all nvidia/cuda nvidia-smi# run docker container without CUDA-toolkit installed
docker run -i --gpus all ubuntu nvidia-smi
```

7. Enable access to S3 Buckets from a Ubuntu 18.04 Amazon Machine Image

Beforehand, an Amazon Identity Access Management (IAM) instance profile role must be created and attach, at least, the AmazonS3FullAccess permissions policy. More policies can be attached if the instances are intended to be used within AWS Batch. (check example screenshot below)

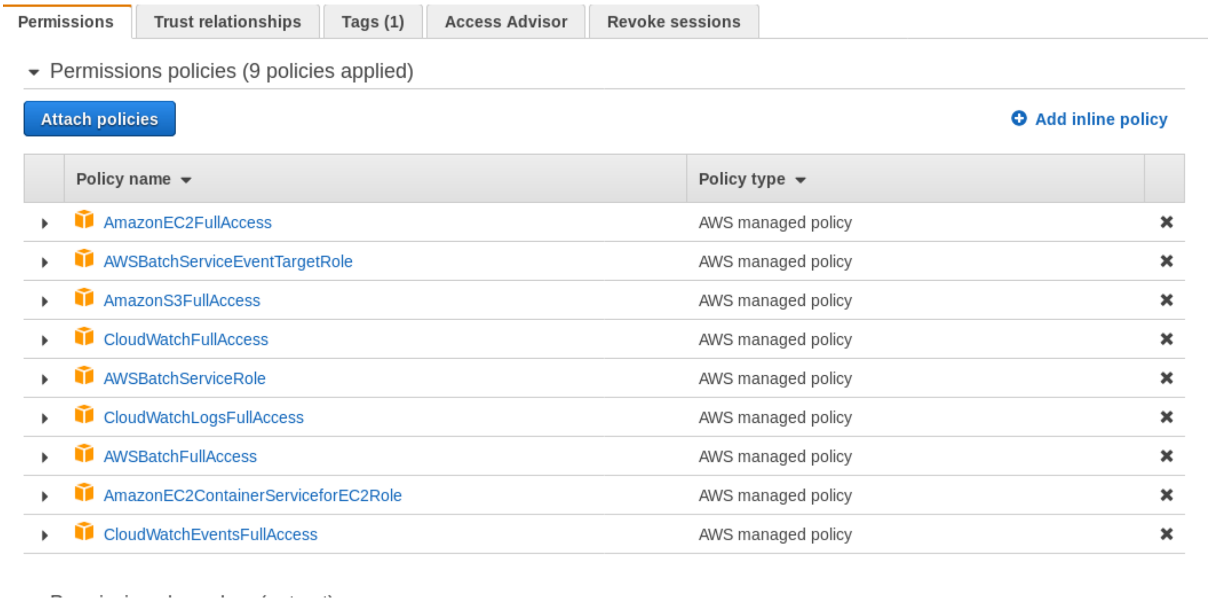


```
# Reference: https://geraldalinio.com/aws/s3/install-s3fs-and-mount-s3-bucket-on-ubuntu-18-04-
instance/
# Install S3fs Fuse drive as root
apt-get install automake autotools-dev fuse g++ git libcurl4-gnutls-dev libfuse-dev libssl-dev
libxml2-dev make pkg-config
git clone https://github.com/s3fs-fuse/s3fs-fuse.git
cd s3fs-fuse
./autogen.sh
./configure
make
make install

echo ACCESS_KEY_ID:SECRET_ACCESS_KEY > /root/.passwd-s3fs
chmod 600 /root/.passwd-s3fs
```

```
#Line in /etc/fstab - Adapt according to your bucket named and preferred mounting point. Modify url parameter depending on the used Amazon computing region. Values uig

frankfurt-nf     /mnt/frankfurt-nf fuse.s3fs _netdev,allow_other,passwd_file=/root/.passwd-s3fs,url=https://s3-eu-central-1.amazonaws.com,uid=1000,gid=1000   0 0
```
